# IL-15 priming alters IFN-γ regulation in murine NK cells

**DOI:** 10.1101/2023.04.23.537947

**Authors:** Maria Cimpean, Molly P. Keppel, Anastasiia Gainullina, Changxu Fan, Nathan C. Schedler, Amanda Swain, Ana Kolicheski, Hannah Shapiro, Howard A. Young, Ting Wang, Maxim N. Artyomov, Megan A. Cooper

**Affiliations:** Department of Pediatrics, Division of Rheumatology/Immunology, Washington University School of Medicine, St. Louis, MO 63110, USA; Department of Pathology and Immunology, Washington University School of Medicine, St. Louis, MO 63110, USA; Department of Genetics, Center for Genome Sciences and Systems Biology, Washington University School of Medicine, St. Louis, MO 63110, USA; Cancer Innovation Laboratory, Center for Cancer Research, National Cancer Institute, Frederick, MD, United States

## Abstract

Natural killer (NK) effector functions can be triggered by inflammatory cytokines and engagement of activating receptors. NK cell production of IFN-γ, an important immunoregulatory cytokine, exhibits activation-specific IFN-γ regulation. Resting murine NK cells exhibit activation-specific metabolic requirements for IFN-γ production, which are reversed for activating receptor-mediated stimulation following IL-15 priming. While both cytokine and activating receptor stimulation leads to similar IFN-γ protein production, only cytokine stimulation upregulates *Ifng* transcript, suggesting that protein production is translationally regulated after receptor stimulation. Based on these differences in IFN-γ regulation, we hypothesized that ex vivo IL-15 priming of murine NK cells allows a switch to IFN-γ transcription upon activating receptor engagement. Transcriptional analysis of primed NK cells compared to naïve cells or cells cultured with low-dose IL-15 demonstrated that primed cells strongly upregulated *Ifng* transcript following activating receptor stimulation. This was not due to chromatin accessibility changes in the *Ifng* locus or changes in ITAM signaling, but was associated with a distinct transcriptional signature induced by ITAM stimulation of primed compared to naïve NK cells. Transcriptional analyses identified a common signature of c-Myc (Myc) targets associated with *Ifng* transcription. While Myc marked NK cells capable of *Ifng* transcription, Myc itself was not required for *Ifng* transcription using a genetic model of Myc deletion. This work highlights altered regulatory networks in IL-15 primed cells, resulting in distinct gene expression patterns and IFN-γ regulation in response to activating receptor stimulation.

## Introduction

Natural killer (NK) cells are innate lymphocytes important for the immune responses to infections and tumors. They are aptly named after their ability to recognize and eliminate infected, “stressed”, or malignant cells through directed release of their cytotoxic granule content or death receptor-mediated apoptosis [1, 2]. NK cells are a major producer of interferon-gamma (IFN-γ) prior to antigen-specific T cell responses, which is important for modulation of the immune response, supporting optimal anti-viral and anti-tumor responses through effects such as enhanced macrophage function and upregulation of antigen presentation-related proteins [3–6].

NK effector functions are controlled by cytokines and the integration of activating and inhibitory signals from germline-encoded receptor interactions with ligands on potential target cells [7]. The common gamma cytokine interleukin (IL)-15 supports NK cell proliferation, survival and in vivo persistence, homeostasis, cytotoxicity and cytokine production [8, 9]. Administration of IL-15 in vivo in murine models enhances anti-tumor activity and anti-viral responses [10–12]. In vitro culture systems have delineated effects of low versus high dose IL-15 in both human [13–15] and murine systems [16, 17]. At low concentrations, IL-15 supports NK survival. High dose IL-15 induces proliferation, acquisition of enhanced cytotoxic capacity [16], and metabolic reprogramming through upregulation of mTOR and glycolysis [18, 19]. Upregulation of mTOR and glycolysis supports optimal responses to MCMV infection, including antigen-specific killing, degranulation, and granzyme B expression [12, 18, 20, 21]. Due to the effect of IL-15 on NK cells, IL-15-based therapies have been developed and are undergoing testing in clinical trials, where they have shown promise in enhancing antitumor responses, with enhanced NK cell persistence and expansion without stimulating regulatory T cells [22–24].

We previously reported that IFN-γ production by freshly isolated (naïve) murine NK cells has distinct metabolic requirements dependent on the route of activation. IFN-γ production following activating receptor stimulation, mediated by ITAM signaling, has a high metabolic requirement, and both glycolysis and mitochondrial oxidative phosphorylation (OXPHOS) are needed. By contrast, cytokine-stimulated IFN-γ production remains intact with metabolic inhibition, demonstrating metabolic flexibility [19]. The strict metabolic requirement for ITAM signaling can be reversed by culturing NK cells with high dose IL-15 for 72 hours (priming) in vitro [19], and in vivo by administration of IL-15 leading to rescue of otherwise fatal infection with murine cytomegalovirus (MCMV) when glycolysis is inhibited [12].

In this study, we characterized the effects of IL-15 priming on NK cells, focusing on the changes in IFN-γ regulation downstream of activating receptor stimulation. We previously demonstrated that while cytokine stimulation induces rapid upregulation of *Ifng* transcript followed by protein, ITAM-based stimulation leads to similar IFN-γ protein production but without upregulation of transcript [19]. Based on these differences in metabolic requirements and *Ifng* expression, we hypothesized that transcriptional regulation of *Ifng* is related to the metabolic requirement for protein production, with new transcription/translation having metabolic flexibility. Here, we identify that IL-15 priming alters chromatin accessibility and leads to transcriptional regulation of IFN-γ production in response to ITAM-signaling, fundamentally altering the signals received by NK cells through activating receptors. Acquisition of metabolic flexibility is associated with induction of c-Myc, which marks cells capable of *Ifng* transcription, but is not itself required for transcription. Overall, this study provides insight into mechanisms of IL-15 priming and has implications for the design of therapeutic approaches to optimize NK cell effector functions in metabolically challenging environments in vivo [21, 25].

## Materials and Methods

### Mice and NK cell enrichment

All mice were maintained in specific pathogen-free conditions and studies were approved by the Washington University Animal Studies Committee. Wild-type mice were purchased from Charles River (C57BL/6NCrl, #556) and used for studies between 6 and 14 weeks of age. Splenic NK cells were enriched using a negative selection magnetic bead kit (STEMCELL Technologies). NK cell purity was 95% or higher, and the majority of contaminating cells were CD3 negative. Unless otherwise specified, controls were C57BL/6NCrl. Heterozygous ΔARE mice were generated and obtained from Howard A. Young’s lab and bred in our animal facility with WT mice. These mice have a targeted substitution of a conserved 162 nucleotide AU-rich sequence in the 3’ untranslated region (3’UTR) of the IFN-γ [26].

Reporter mice were obtained from The Jackson Laboratory. IFN-γ YFP reporter mice on a C57BL/6 background (B6.129S4-Ifngtm3.1Lky/J, JAX stock #017581) have an IRES-eYFP reporter cassette inserted between the translational stop codon and 3’ UTR/polyA tail of the interferon gamma (*Ifng*) gene (referred to as GREAT mice [27]). Myc reporter mice (Myc^EGFP^) (B6;129-Myctm1Slek/J, JAX stock #021935) express an N-terminal fusion EGFP/MYC protein [28]. Myc^EGFP^ mice with NK1.1 expression comparable to a C57BL/6 background were used for experiments.

An NK cell and ILC1-specific inducible model of c-Myc deletion, *Ncr1*-*Myc*Δ/Δ, was generated by crossing mice with yellow fluorescent protein (YFP) Cre-reporter and tamoxifen-inducible iCre recombinase driven by *Ncr1* (*Ncr1*-iCreERT2) [29] with mice carrying loxP sites flanking exons 2 and 3 of c-Myc [30] (generated by the Alt lab [30] and generously shared by Dr. Ruoning Wang at Nationwide Children’s Hospital). N*cr1*-iCreERT2 heterozygous mice (*Ncr1*-WT) were used as controls for experiments where Myc deletion was induced with tamoxifen chow or 4-OHT. The controls were housed in the same facility but were not littermates or co-housed.

### Cell culture

Complete media (“R10”) contained RPMI 1640 (Corning), 10% fetal bovine serum (FBS) (Sigma Aldrich), L-glutamine (Sigma Aldrich), non-essential amino acids (Corning), penicillin/streptomycin (Sigma Aldrich). For glucose-free media, the following substitutions/replacements were made: glucose-free RPMI (Corning), 15% PBS (Corning), and 10% dialyzed FBS. For 72 hour cultures, media also contained 50 μM β-mercaptoethanol (Sigma Aldrich) and 10 ng/ml (LD) or 100 ng/ml (HD) IL-15 (Peprotech). For experiments with 4-hydroxytamoxifen (4-OHT), 0.8 uM was the final 4-OHT concentration in vitro. Metabolic or pathway inhibitors were used at the following final concentrations: 2DG (1mM, Sigma Aldrich), rapamycin (10 nM, 100 nM), oligomycin (10 or 100 nM).

### Human NK cells

Cryopreserved human peripheral blood mononuclear cells (PBMCs) from anonymous healthy platelet donors (de-identified Leukoreduction System chambers, non-human subjects) were thawed and rested overnight in complete media (10% human AB serum). Cells were then washed with PBS and resuspended in either complete or glucose-free media. The cells were stimulated with IL-12+IL-18 (10 ng/ml, Peprotech, and 50 ng/ml, MBL, respectively) or plate-bound anti-CD16 (1 mg/ml) for 6 hours.

### Flow cytometry

A live/dead stain was included in most assays to measure viability (Zombie Yellow, BioLegend). After murine NK cell enrichment, which consistently resulted in a ∼95% CD3^-^NK1.1^+^ or CD3^-^ NKp46^+^ population, cells were gated on NKp46 to identify NK cells. For human NK cells, cells were gated based on Zombie Yellow to identify live cells, followed by CD3 and CD56 to identify NK cells. Murine cells were blocked with anti-FcγRIII (2.4G2) prior to surface staining. Human cells were blocked with mouse IgG prior to surface staining. Cells were fixed with 1% paraformaldehyde for surface analysis or fixed with Cytofix/Cytoperm (BD Bioscience) for intracellular staining. For phospho flow assays, cells were fixed with 2% paraformaldehyde, permeabilized with methanol, and stained overnight at 4C with phospho antibodies. Data was acquired on a Cytek-modified FACScan (BD and Cytek) or LSRFortessa (BD); data was analyzed using FlowJo software. The following fluorochrome-conjugated antibodies were used: CD3ε (145-2C11, BD/BioLegend), CD19 (1D3, BioLegend), CD56 (NKH-1-RD1, Beckman Coulter), ERK1/2 (T202/Y204) (6B8B69, BD), anti-human IFN-γ (B27, BD), anti-mouse IFN-γ (XMG1.2, BioLegend), NFkB p65 (S536) (93H1, BD), NK1.1 (PK136, BD), NKp46 (29A1.4, BD), p38 MAPK (T180/Y182) (36/p38, BD), PLCg2 (Y759) (K86-689.37, BD), ZAP70 (Y319)/Syk(Y352) (XMG1.2, BD).

### NK cell stimulation and IFN-γ production

For cytokine stimulation, NK cells were cultured for 6 hours in the presence of 1 ng/ml murine IL-12 (Peprotech) and 1 ng/ml murine IL-18 (MBL). NK1.1 was engaged by culturing NK cells in plates coated with 20 μg/ml purified anti-NK1.1 (PK136; BioXcell). Where indicated, PMA (Sigma Aldrich) and calcimycin (Sigma Aldrich) were used at 10 ng/ml and 500 ng/ml, respectively. For experiments with actinomycin, freshly isolated splenic NK cells were stimulated in the presence of the indicated doses of actinomycin (Sigma Aldrich). For phospho-flow experiments, cells were incubated with anti-NK1.1 (PK136) and anti-NKp46 (29A1.4) for 10-15 min, followed by the addition of goat anti-mouse antibody (BioLegend) to begin the NK1.1 stimulation. For assays measuring intracellular IFN-γ production, GolgiPlug^TM^ (BD Biosciences, Brefeldin A) was added after 1 h of stimulation in order to inhibit IFN-γ secretion. BD Cytofix/Cytoperm^TM^ (BD Biosciences) was used as recommended for intracellular staining.

### Quantitative PCR

NK cells (∼2-3x10^5) were activated with plate-bound anti-NK1.1 (20 μg/ml) and harvested at the indicated time points. RNA was isolated with the RNeasy Plus Micro kit (Qiagen) and cDNA was generated using random hexamers (Promega), following the GoScript Reverse Transcription System protocol (Promega). Primers for *Actb* (Mm00607939, Thermo Fisher Scientific), *Ifng* ([31], IDT) and *Myc* (Mm00487804_m1, Thermo Fisher Scientific) were used to evaluate gene expression. Copy numbers of *Ifng* and *Actb* transcript were determined by standard curves generated with plasmid clones of *Ifng* and *Actb* amplicons and quantified via real-time quantitative PCR (TaqMan, QuantStudio 3 instrument from Thermo Fischer Scientific). To normalize across conditions, *Ifng* copy number relative to *Actb* was calculated. For *Myc* expression, data were analyzed using the 2^−ΔΔCt^ method [32].

### ATAC-seq

Aliquots of approximately 75,000 cells (naïve, LD/HD IL-15) were used as input and processed as previously described [33]. Briefly, the cells were washed in cold PBS and lysed in cold lysis buffer (10mM Tris-HCl pH 7.4, 10mM NaCl, 3 mM MgCl2, 0.1 % IGEPAL CA-630). Transposase reaction occurred at 37C for 30 minutes. DNA was purified with the Qiagen MinElute PCR purification kit and amplified for 5 cycles. The appropriate number of additional PCR cycles was determined by real-time PCR. The final product was cleaned using AMPure XP beads (Beckman Coulter). Quality control of the generated libraries was performed using the Agilent TapeStation system. Libraries were sequenced on a NovaSeq6000.

ATAC-seq data were first processed using the AIAP [34] pipeline. Briefly, reads were aligned to the mm10 genome using bwa; locations of Tn5 insertion events were inferred from properly paired, non-PCR duplicate reads with MAPQ >= 10; each Tn5 insertion event was extended from insertion site up and down 75 bp to generate a 150 bp “pseudo read”; pseudo reads were piled up to generate an ATAC signal profile, which was subsequently normalized against both sequencing depth and sample quality, similar to the getGroupBW() function of the ArchR package [35]. Briefly, sample quality was calculated as the fraction of reads mapped to promoters (transcription start sites ± 1 Kb. frac.pro); a sequencing depth scaling factor (depth.fac) was calculated for each sample as the ratio of the sequencing depth of that sample over the median sequencing depth of all samples; each ATAC-seq profile (bigWig track) was then divided by frac.pro and depth.fac to obtain normalized signals; normalized ATAC-seq signals were divided by 3 because the typical value of depth.fac is 0.3 - 0.4. ATAC-seq data were visualized using WashU Epigenome Browser [36].

ATAC-seq peaks for each sample were called as a part of the AIAP pipeline, using macs2 [37] with FDR cutoff at 0.01. After manual inspection of the peaks called, we further filtered the resulting narrowPeak files for peaks with -log10FDR >= 8. To call differentially accessible regions (DARs), we merged all overlapping peaks to create a union peakset. We then formed a peak-sample matrix with each entry corresponding to the number of Tn5 insertion sites in each peak for each sample. We filtered out peaks where any sample has no reads. DARs were called using DESeq2 [38] (v1.26.0), with the design formula ∼ replicate + type. “Type” represents high or low dose IL15 treatment. The cutoff for DARs were set at log2FoldChange > 1 & FDR < 0.05.

For motif enrichment in DARs, up (high dose > low dose) and down (low dose > high dose) DARs were separately ranked by p value and the top 1000 peaks in each category were used for motif enrichment. For each DAR used for motif enrichment, its 25 closest non-DARs in the Euclidean space constructed from low dose samples were selected without replacement as background peaks. We used HOMER (v4.11.1) [39] for motif enrichment in DARs relative to background peaks (findMotifsGenome.pl up_or_down_DAR.bed mm10.fa out_dir -bg up_or_down_DAR_background.bed -nomotif -size given).

### RNA-seq analysis

Enriched NK cells (>95% purity) were either cultured for 72 hours in media containing LD or HD IL-15, or used immediately after enrichment. Where indicated, cells were stimulated with anti-NK1.1 or IL-12 and IL-18 as described above. RNA was isolated using Dynabeads mRNA DIRECT Kit (Invitrogen), and then reverse transcribed using the AffinityScript Multiple Temperature cDNA Synthesis Kit (Agilent Technologies) to synthesize cDNA. After first strand synthesis, samples were pooled together based on ACTB qPCR values, and RNA-DNA hybrids degraded using consecutive acid-alkali treatment. A second sequencing linker (AGATCGGAAGAGCACACGTCTG) was ligated using T4 ligase (NEB) followed by SPRI clean-up. The mixtures were then PCR enriched for 14 (IL-15 dataset) or 17-20 (activation dataset) cycles and SPRI purified to yield final strand specific RNA-seq libraries. Final library QC on pooled libraries was performed using the Agilent TapeStation system. Libraries were sequenced on HiSeq 2500 (Illumina) using 40 bp X 10 bp paired-end sequencing. Second mate was used for sample demultiplexing, generating individual single-end fastq files.

Reads were aligned to the mouse genome GRCm38/mm10 primary assembly (GENCODE) and gene annotation Ver.M16 with STAR 2.5.4a. The raw read counts were generated by featureCounts [40] and normalized with DESeq2 [38] package from R/Bioconductor. The top 12,000 genes ranked by average gene expression were selected for differential expression analysis using the DESeq2 [38]. The cutoff for determining significantly differentially expressed genes was an FDR-adjusted p value less than 0.05. Differential expression analysis was performed with the limma package via Phantasus, a web-based application [41]. Bubble plots and GSEA [42] plots were generated in R with the enrichplot package [43].

For the IFN-γ reporter (GREAT) dataset, cells were primed as above and stimulated via plate-bound anti-NK1.1 for 6 hours, then sorted on a FACSAria Fusion (BD Biosciences) based on YFP. RNeasy Plus Micro kit (Qiagen) was used for RNA extraction. Samples were prepared according to the library kit manufacturer’s protocol, indexed, pooled, and sequenced on an Illumina NovaSeq 6000. Basecalls and demultiplexing were performed with Illumina’s bcl2fastq software and a custom python demultiplexing program with a maximum of one mismatch in the indexing read. RNA-seq reads were then aligned to the Ensembl release 76 primary assembly with STAR version 2.7.9a [44]. Gene counts were derived from the number of uniquely aligned unambiguous reads by Subread:featureCount version 2.0.3 [40]. Sequencing performance was assessed for the total number of aligned reads, total number of uniquely aligned reads, and features detected. The ribosomal fraction, known junction saturation, and read distribution over known gene models were quantified with RSeQC version 4.0 [45].

All gene counts were then imported into the R/Bioconductor package EdgeR [46] and TMM normalization size factors were calculated to adjust for samples for differences in library size. Ribosomal genes and genes not expressed in the smallest group size minus one samples greater than one count-per-million were excluded from further analysis. The TMM size factors and the matrix of counts were then imported into the R/Bioconductor package Limma [47]. Weighted likelihoods based on the observed mean-variance relationship of every gene and sample were then calculated for all samples and the count matrix was transformed to moderated log 2 counts-per-million with Limma’s voomWithQualityWeights [48]. The performance of all genes was assessed with plots of the residual standard deviation of every gene to their average log-count with a robustly fitted trend line of the residuals. Differential expression analysis was then performed to analyze for differences between conditions and the results were filtered for only those genes with Benjamini-Hochberg false-discovery rate adjusted p-values less than or equal to 0.05.

## Data Availability

RNA and ATAC sequencing data generated during this study is available at Gene Expression Omnibus with accession number GSE227894.

## Results

### Regulation of IFN-γ in naïve and IL-15 primed NK cells

We previously demonstrated that IL-12+IL-18 activation of IFN-γ protein production is transcriptionally regulated in freshly isolated (naïve) murine NK cells, whereas ITAM-based (anti-NK1.1) activation leads to similar protein production, but not IFN-γ transcript [19]. To investigate how IFN-γ protein production is regulated by cytokine or activating receptor stimulation, we first assessed the contribution of transcription to IFN-γ production after IL-12+IL-18 or plate-bound anti-NK1.1 stimulation by culturing the cells in the presence of the transcription inhibitor actinomycin D. While the percentage of IFN-γ+ NK cells decreased with both stimulations, the decrease was significantly more pronounced in the cytokine stimulated cells (**Fig. 1A**). This supports the hypothesis that IFN-γ is predominantly translated from pre-formed transcript when NK cells are activated via ITAM-bearing receptors, while cytokine (IL-12/18) stimulation is highly dependent on new transcription.

**Figure 1.**
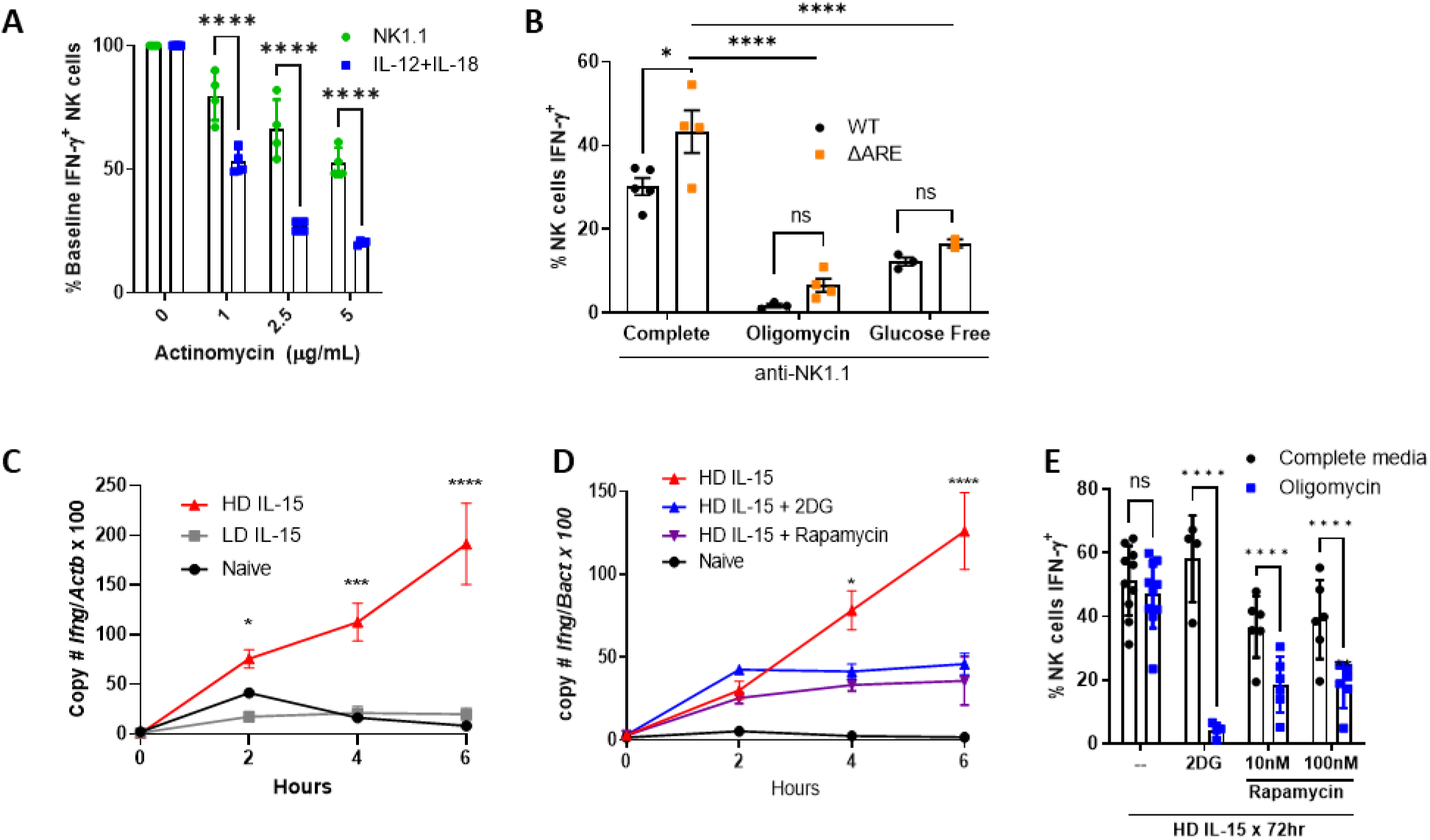
Regulation of IFN-γ production in naïve and IL-15 primed NK cells. **A**) Freshly isolated NK cells stimulated with plate-bound anti-NK1.1 (PK136) or low dose IL-12+IL-18 (1 ng/ml each) in the presence of indicated doses of actinomycin. % maximal IFN-γ+ NK cells shown compared to baseline (no actinomycin). 2 independent experiment, duplicate wells; **B**) NK cells from WT mice or mice with deleted 3’ AU rich elements (ARE) of Ifng were cultured in complete media +/- 100nM oligomycin or in glucose-free media during the 6 hour anti-NK1.1 stimulation. n=4-5 mice per genotype, 2 independent experiments. **C-E**) NK cells were freshly isolated (naïve) or cultured for 72 hours with 10 ng/ml (LD) or 100 ng/ml (HD) IL-15, followed by anti-NK1.1 stimulation for the indicated time, up to 6 hours. **C**) Absolute copy number of Ifng and Actb (beta-actin) were quantified by real-time PCR. Results represent the ratio of Ifng to Actb x 100 and the mean +/- SEM of duplicate wells from 2 independent experiments. **D-E**) NK cells were primed in HD IL-15 in the absence or presence of 2DG (1 mM) or rapamycin (100 nM) prior to NK1.1 stimulation. **D**) Ratio of Ifng to Actb x 100 and the mean +/- SEM from 1-3 independent experiments. **E**) % IFN-γ+ NK cells were measured by flow cytometry after 6h anti-NK1.1 stimulation, 2-6 independent experiments, 4-10 mice per condition. Statistical analysis: 2way ANOVA (**A**, **C-E**), Ordinary one-way ANOVA (**B**).

In addition to differences in transcriptional or post-transcriptional regulation of IFN-γ production, we also previously demonstrated that there are distinct metabolic requirements for IFN-γ production. Activating receptor-stimulated IFN-γ production is highly sensitive to metabolic inhibitors of both OXPHOS and glycolysis. In contrast, IL-12+IL-18 cytokine-stimulated IFN-γ protein production, is metabolism-independent [19]. This dependence on a secondary metabolic signal for IFN-γ production downstream of ITAM signaling was also observed in human NK cells (**Fig. S1)**. However, we previously demonstrated that culturing murine NK cells with high dose (HD, 100ng/ml) IL-15 for 72 hours (IL-15 priming) leads to metabolic flexibility upon activating receptor stimulation, with intact IFN-γ production in the presence of metabolic inhibitors, similar to IL-12+IL-18 stimulated naive NK cells [19]. The mechanism of this metabolism-independent receptor-stimulated IFN-γ production is relevant when considering how to prepare NK cells for metabolically challenging microenvironments, for example with NK cell adoptive immunotherapy [25].

One potential mechanism for metabolic regulation of IFN-γ protein production may be through GAPDH acting as an IFN-γ translation repressor as demonstrated in CD4+ T cells. In T cells, GAPDH binds to AU-rich elements (ARE) in the 3’ untranslated region (UTR) of IFN-γ mRNA [49]. Upon glycolysis upregulation, which engages GAPDH enzymatic activity, IFN-γ translation is no longer inhibited in CD4+ T cells. To test if a 3’UTR-dependent mechanism also occurs in naïve NK cells, we evaluated the effect of metabolic inhibition on IFN-γ production in mice with a 162 nucleotide ARE region substitution in the 3’UTR of the IFN-γ gene (ΔARE) [50]. NK cells in ΔARE mice have been shown to have elevated IFN-γ protein production at baseline [26], and we observed this with receptor stimulation (**Fig. 1B** complete). In this genetic model, metabolic inhibitors, 2-DG to inhibit glycolysis or oligomycin to inhibit mitochondrial OXPHOS, both inhibited activating-receptor induced IFN-γ production similar to wild-type (WT), suggesting that the mechanism of metabolic dependence for activating receptor stimulation in NK cells is 3’UTR-independent (**Fig. 1B**).

Based on the observed differences in IFN-γ regulation, we hypothesized that IL-15 priming of NK cells allows a switch to IFN-γ transcriptional regulation upon receptor activation, thus allowing for metabolic flexibility. To determine the effect of IL-15 priming on NK1.1-mediated *Ifng* transcription, *Ifng* copy number was measured in anti-NK1.1 stimulated NK cells that were either freshly isolated (naïve) or primed for 72 hours with a high-dose IL-15 (HD, 100 ng/ml - primed cells) or cultured in the presence of low-dose (LD, 10 ng/ml), which does not lead to metabolic flexibility [19]. At baseline (time 0), *Ifng* copy number by qPCR was similar across conditions, demonstrating that IL-15 priming itself does not upregulate *Ifng* transcript. Over the course of the 6 hour anti-NK1.1 stimulation, *Ifng* significantly increased only in NK cells previously cultured in HD IL-15. By contrast, cells cultured in LD IL-15, similar to naïve NK cells, expressed very low *Ifng* transcript levels that did not change over the course of the 6 hour anti-NK1.1 stimulation (**Fig. 1C**). These results demonstrate that IL-15 primed cells switch to transcriptional regulation of IFN-γ and that transcriptional regulation of IFN-γ is dependent on IL-15 dose and associated with metabolic flexibility.

Culturing NK cells in HD IL-15 promotes proliferation and increases metabolic activity with a prominent upregulation of glycolysis [18, 19]. Undivided cells produce similar amounts of IFN-γ protein after HD IL-15 priming, suggesting that the change in metabolic requirements is not related to proliferation [19]. High dose IL-15 activates mTOR and results in increased glucose uptake [18]. In short term culture experiments, IL-15-induced mTOR is important for NK IFN-γ production [51]. To evaluate the contribution of glycolysis and mTOR to the IL-15 priming effect on IFN-γ transcription, NK cells were primed with HD-IL-15 in the presence of 2-DG or rapamycin followed by activating receptor stimulation (anti-NK1.1). Cells primed in the presence of these inhibitors had impaired upregulation of *Ifng* transcript production, with reduced protein production that was most prominent in cells cultured with 2DG (**Fig. 1D, 1E**). This suggests that the ability to upregulate glycolysis, and mTORC1, is critical during the priming phase for the IL-15 priming effect.

### IL-15 priming induces chromatin accessibility changes

To study the effect of IL-15 priming on the epigenome and regulatory network of NK cells, ATAC-seq was performed on primed NK cells. Differentially accessible chromatin regions were compared with NK cells cultured with LD IL-15, which do not upregulate *Ifng* transcript and lack metabolic flexibility, to account for changes associated with in vitro culture. There were 5120 differentially accessible regions (DARs), 3845 of which were more accessible and 1275 of which were less accessible in primed NK cells. For example, one of the genes with several regions of increased chromatin accessibility was *Spp1* (**Fig. 2A, 2C**). *Spp1* encodes for secreted phosphoprotein 1 (osteopontin), which is upregulated downstream of IL-15 signaling [52]. The more accessible (HD>LD) or “up” DARs displayed a strong enrichment for motifs bound by transcription factors Fos, JunB, Fra1, Fra2, and BATF, sharing a common binding sequence (5’ - TGA(C/G)TCA - 3’) (**Fig. 2B**). With these transcription factors representing members of the AP-1 transcription factor family, the results suggest that IL-15 primed NK cells may be more responsive to activation of the MAPK pathway, and subsequent activation of AP-1 family members.

**Figure 2.**
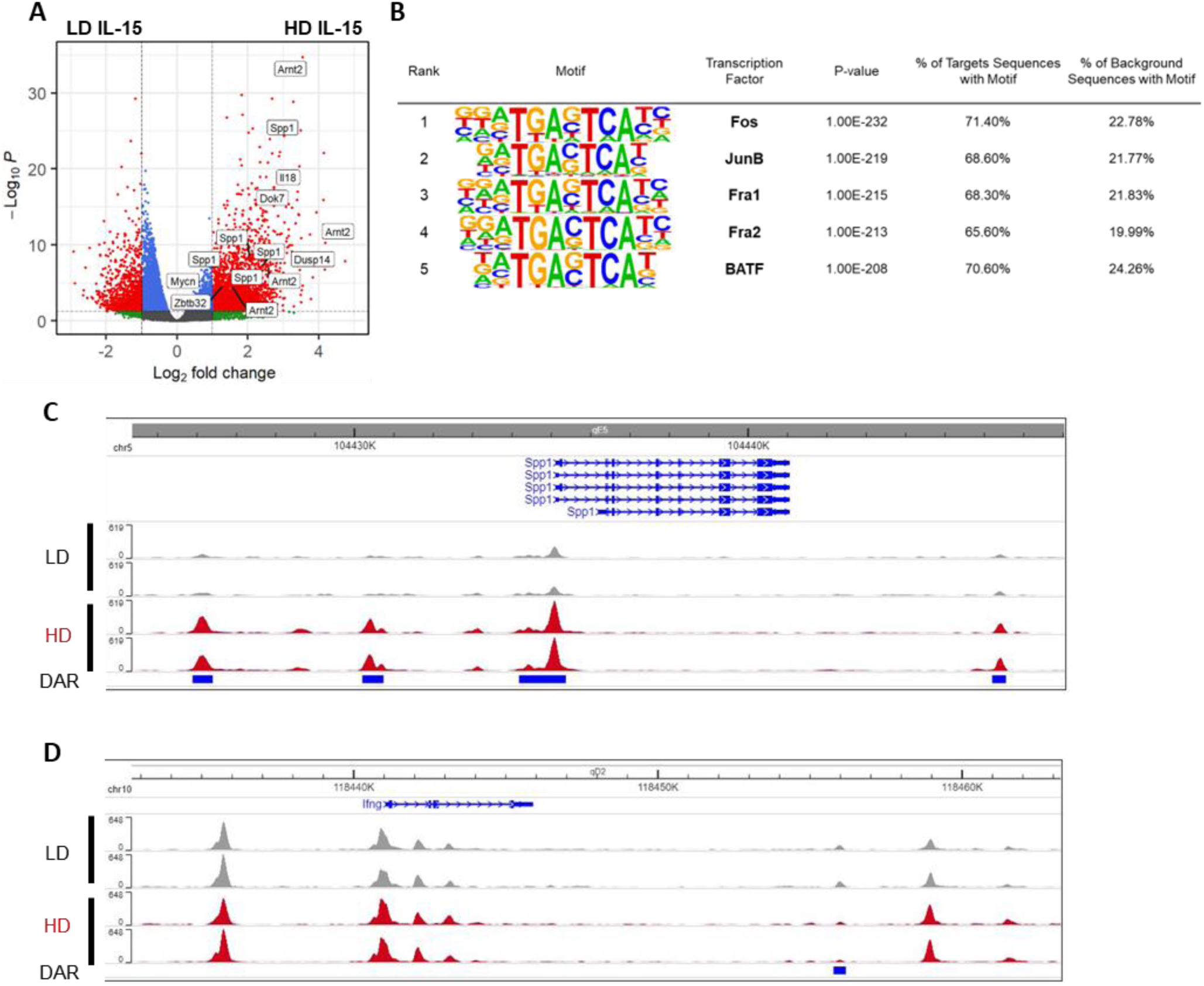
IL-15 priming induces chromatin accessibility changes. ATAC-seq analysis comparing LD vs HD IL-15 cultured NK cells, after 72 hours. 2 mice per group. **A**) Volcano plot showing the differentially accessible (log2FoldChange > 1, FDR < 0.05) chromatin regions assigned to genes. **B**) Motif enrichment at regions of open chromatin as defined by ATAC-seq in primed NK cells compared to NK cells cultured in LD IL-15. **C-D**) Genome browser visualization of peaks near *Spp1* (C) and *Ifng* (D).

Given the time-dependent effect of IL-15 priming on *Ifng* regulation, we hypothesized that priming might induce chromatin accessibility changes in the *Ifng* locus, including conserved non-coding sequences that contribute regulatory functions to the *Ifng* locus, potentially creating new sites for transcription factors to occupy and modulate *Ifng* transcription. However, the *Ifng* region remains almost unchanged, with the exception of one DAR located approximately 15kb from the TSS (**Fig. 2D**). This DAR is modestly less accessible in IL-15 primed NK cells and has an undetermined biological significance. Therefore, chromatin accessibility changes, directly at regulatory elements in the *Ifng* locus, are unlikely to play a part in the switch to *Ifng* transcriptional regulation. Collectively, this data demonstrates that culturing NK cells in high dose IL-15 for 72 hours predominantly increases chromatin accessibility across the genome in murine NK cells, but not in the *Ifng* locus.

### IL-15 priming does not alter canonical ITAM signaling in a dose-dependent manner

With the knowledge that there are no up DARs in the *Ifng* locus induced by IL-15 priming, an alternative hypothesis for transcriptional regulation of *Ifng* with priming was that strength of ITAM signaling is altered in by IL-15 priming. Based on current knowledge of ITAM-based signaling in NK cells (**Fig. 3A**) [7, 53], phosphorylation of relevant proximal and distal signaling molecules including Zap70, Syk, PLCγ2, MAPK Erk and p38, and p65 (RelA) following anti-NK1.1 stimulation was measured by flow cytometry (**Fig. 3B-E**). There were no differences in the mean fluorescence intensity (MFI) of these phosphorylated proteins in NK cells cultured with LD IL-15 versus HD IL-15 (primed) (**Fig. 3B-E**). Similarly, the presence of oligomycin or culture in glucose-free media during the stimulation did not alter ITAM signaling (**Fig. S2**). These results demonstrate that there are no significant differences in canonical ITAM signaling that correlate with the switch to transcriptional regulation *Ifng* after IL-15 priming.

**Figure 3.**
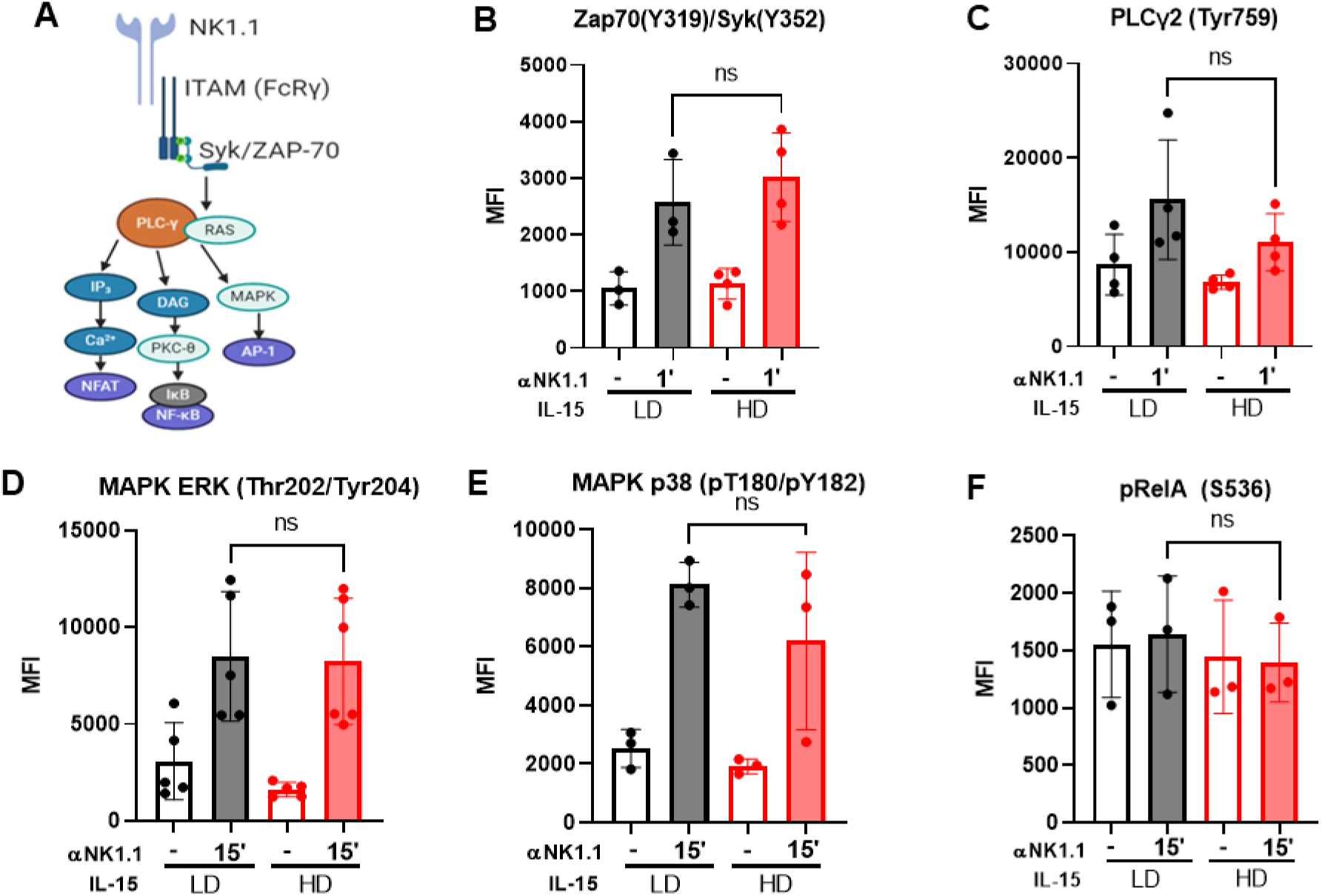
IL-15 does not alter ITAM signaling in a dose-dependent manner. **A**) ITAM signaling pathways downstream of NK1.1. **B-F**) Murine NK cells were cultured for 72 hours in LD (10 ng/ml) or HD (100 ng/ml) IL-15. Cells were washed, then cultured for 30-60 min in complete media (+/- 100 nM oligomycin) or glucose-free media. Phosphorylation of Zap70/SYK (**B**), PLCγ2 (**C**), ERK (**D**), MAPK p38 (**E**), and RelA (**F**) was quantified by flow cytometry. Results shown are from a minimum of 3 independent experiments, 1 mouse per experiment. Statistical analysis: Wilcoxon test (paired t-test, non-parametric)

### Transcriptional changes associated with *Ifng* transcription in IL-15 primed NK cells

While ITAM signaling itself appears unchanged, the regulation of IFN-γ is different in primed NK cells. Given that chromatin accessibility changes, we hypothesized that altered chromatin accessibility in primed NK cells leads to a differential transcriptional response in how NK cells receive those signals. To determine if primed NK cells have a different transcriptional response to activating receptor stimulation, RNA-seq was performed to compare the responses of naïve, LD IL-15 and primed NK cells (**Fig. 4A**). First, to examine transcriptional changes associated with IL-15 priming, unstimulated naïve and HD IL-15 treated NK cells were compared. As expected, and consistent with the reported biology of IL-15 and NK cells, primed NK cells upregulated genes and pathways associated with proliferation, glycolysis, and cytotoxicity [16, 18] (**Fig. 4B**). Glycolysis (*Aldoc*, *Pgam1*, *Eno1*, *Pkm*, *Ldha*; log2FC>2) and cell cycle (*Aurka*, *Ube2c*, *Cdkn3*, *Cdk1*; log2FC>3) genes were upregulated, as well as several granzyme genes, with *Gzmd*, *Gzme*, and *Gzmc* representing the highest fold changes (log2FC>9) (**Fig. 4C**).

**Fig. 4.**
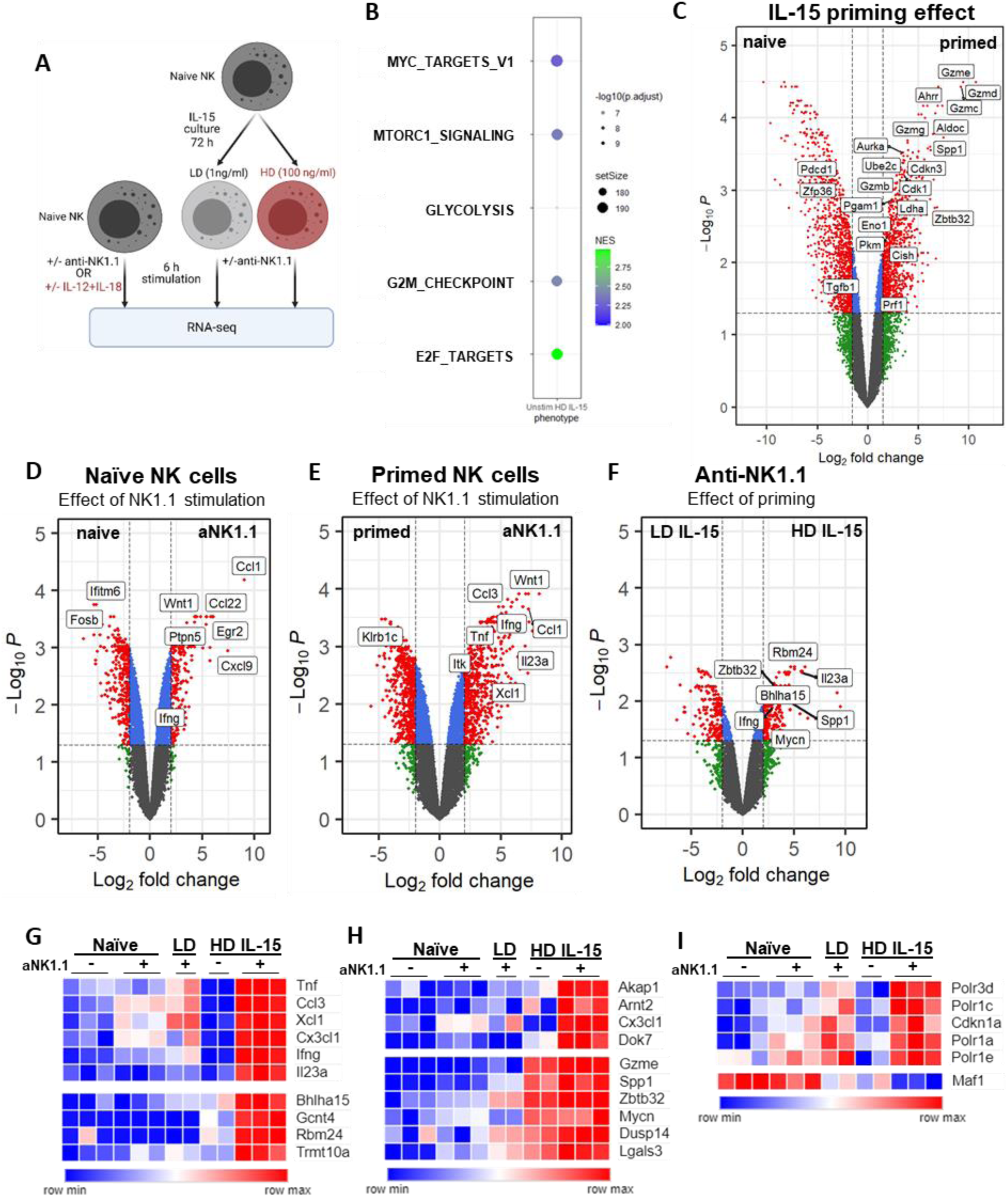
Transcriptional changes associated with *Ifng* transcription. **A**) Experiment schema. NK cells from the spleens of 15 mice were pooled for each dataset (naïve NK +/-stimulation, naïve and IL-15 cultured NK cells +/- anti-NK1.1). Red text indicates conditions with metabolic flexibility for IFN-γ production. **B**) Bubbleplot of gene se enrichment analysis (GSEA) select pathways upregulated in HD unstim compared to naive unstim. **C**) Volcano plot showing differentially expressed genes from the naïve unstim vs HD IL-15 unstim. **D-F**) Volcano plots showing differentially expressed genes from the following comparisons: (**D**) naïve unstimulated vs aNK1.1, (**E**) HD IL-15 unstimulated vs aNK1.1, and (**F**) LD IL-15 aNK1.1 vs HD IL-15 aNK1.1. **G-I**) Heatmap highlighting select genes that are differentially regulated in (**G**) HD+aNK1.1, (**H**) have both increased chromatin accessibility and gene expression in primed NK cells +/- aNK1.1, and (**I**) select Myc targets differentially regulated in HD IL-15+aNK1.1.

Next, NK cell responses to ITAM signaling via anti-NK1.1 in the naïve state versus primed state were evaluated. NK1.1 stimulation of naïve cells strongly upregulated chemokine genes such as *Ccl1* and *Ccl22* rather than *Ifng* (**Fig. 4D**). In contrast, NK1.1 stimulation of primed cells strongly upregulated cytokine genes such as *Ifng* and *Tnf* (**Fig. 4E**). The distinct transcriptional response of primed NK cells to ITAM stimulation was not solely due to in vitro culture, as primed cells and LD IL-15 cultured NK cells also exhibited distinct gene expression patterns after stimulation (**Fig. 4F**). Consistent with our prior data (**Fig. 1C**), *Ifng* transcript was highly upregulated only in NK1.1-activated primed NK cells (**Fig. 4D-G**).

Based on the hypothesis that ITAM signaling leads to a distinct transcriptional signature in IL-15 primed NK cells, allowing for metabolic flexibility, k-means clustering was performed, and genes uniquely upregulated in the NK1.1-stimulated primed NK cells were identified (**Fig.S3**, **Supplemental table 1**). The cluster with upregulated genes only in NK1.1-activated primed cells contained *Ifng*, cytokine subunit *Il23a*, and genes associated with protein synthesis (*Gcnt4*, *Polr3d*, *Trmt10a*) and secretion (*Bhlha15*) compared to NK1.1-stimulated LD IL-15 cultured cells (**Fig. 4G**). Several of these uniquely upregulated genes were also identified to have increased chromatin accessibility in the ATAC-seq dataset, including *Akap1*, *Arnt2*, *Cx3cl1*, *Dok7* (**Fig**. **2A**, **4H**). Additionally, some of the genes with increased chromatin accessibility in primed cells also exhibited increased gene expression compared to naïve unstimulated cells, and their expression further increased in receptor-stimulated primed cells (*Dusp14*, *Gzme*, *Lgals3*, *Mycn*, *Spp1*, *Zbtb32*) (**Fig. 4H**)*. Lgals3*, *Spp1*, and *Zbtb32* have been previously reported to play a role in the regulation of various NK cell functions [52, 54–56]. Like *Ifng* expression patterns, genes encoding several cytokines (*Tnf*, *Il23a*) and chemokines (*Ccl3*, *Cx3cl1*, *Xcl1*) were also upregulated only in NK1.1-activated primed NK cells, although in a different cluster than *Ifng* (**Fig. 4G**). Collectively, these data support that altered chromatin accessibility leads to a differential transcriptional response to ITAM signaling by IL-15 primed NK cells.

Among the genes uniquely upregulated in the NK1.1-stimulated primed NK cells were several Myc targets, including genes encoding for RNA polymerase I and III subunits [57, 58]. *Polr3d* and *Polr1a*,*c*,*e*, to a lesser extent, were uniquely upregulated in NK1.1-stimulated primed cells (**Fig. 4I**). Additionally, *Maf1*, a known negative regulator of polymerase III-mediated transcription [59], was uniquely downregulated. *Cdkn1a*, which Maf1 has been reported to repress through a Pol III-dependent mechanism [60], is also uniquely upregulated in NK1.1-stimulated primed cells (**Fig. 4I**). Thus, IL-15 primed NK cells exhibit unique and distinct transcriptional changes in response to NK1.1 activation.

With the knowledge that *Ifng* is transcriptionally regulated with both IL12/18 and primed NK cells with ITAM stimulation, we hypothesized that these conditions might share a common pathway of transcriptional regulation. Among the 159 genes that overlapped between those conditions, we again identified *Ifng*, *Myc*, and several known *Myc* targets (*Polr3d*, *Polr1e*, *Fosl1*, *Mybbp1a*, etc.) (**Fig. 5A, Supplemental table 2**).

**Figure 5.**
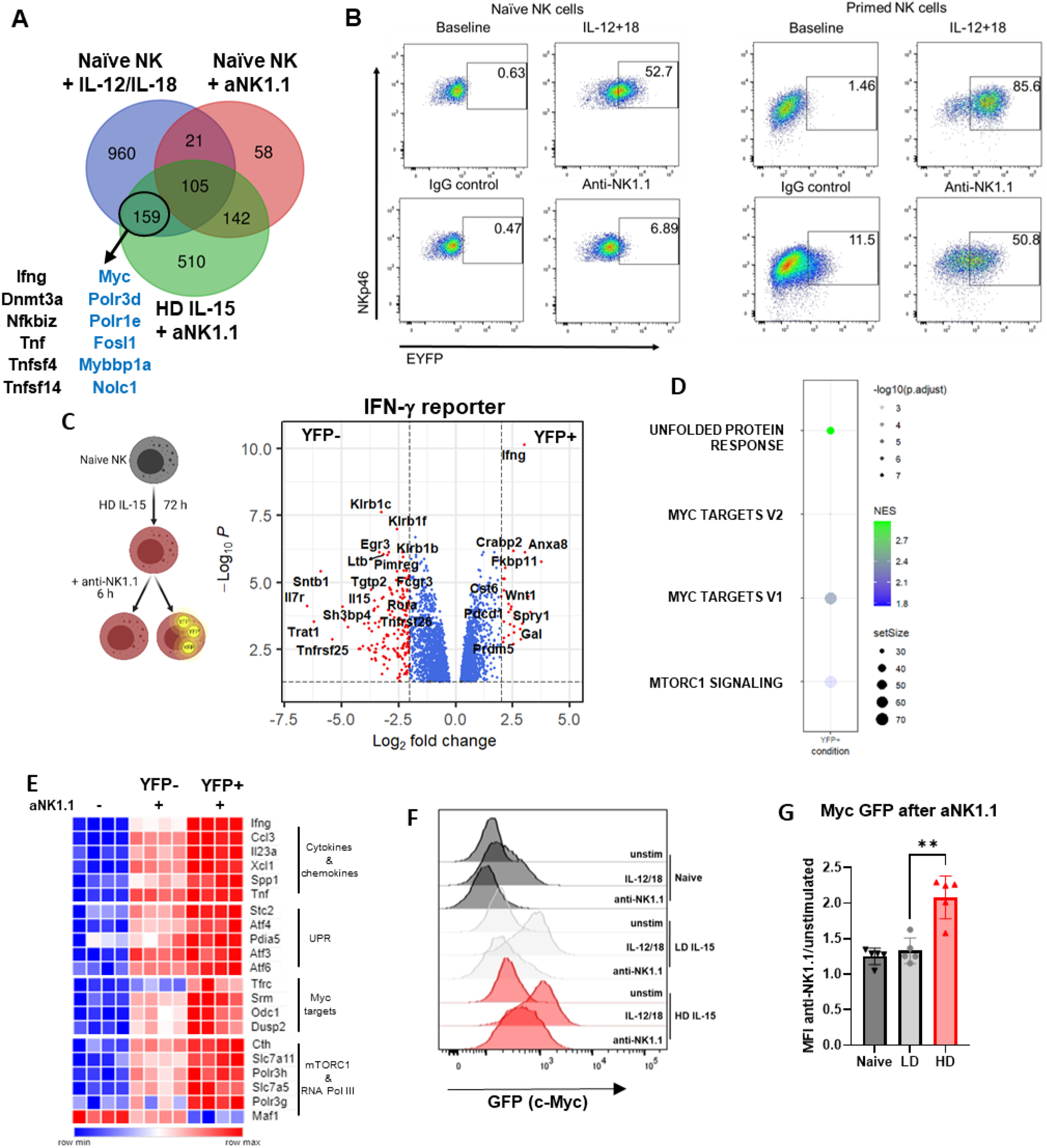
Myc marks NK cells capable of Ifng transcription. **A**) Comparison of genes upregulated in naïve NK cells with IL-12/IL-18 (blue), anti-NK1.1 (red), or in HD IL-15 primed NK cells with anti-NK1.1 (green, compared to HD IL-15 alone). Blue text marks Myc targets. (although *Dnmt3a* has been shown experimentally to be a Myc target). **B**) Representative flow cytometry plots from naïve or primed NK cells from *Ifng* reporter mice, +/- 6 h stimulation. **C**-**D**) NK cells from 5 GREAT mice (3 males, 2 pooled females) were primed with IL-15, then stimulated via plate-bound anti-NK1.1 for 6 hours. Cells were then sorted based on YFP. **C**) Volcano plot shows differentially expressed genes from the IFN-γ+ (YFP+) and IFN-γ- (YFP-) comparison. **D**) Bubbleplot of GSEA analysis pathways upreg in IFN-γ+ compared to IFN-γ-. **E**) Heatmap highlighting select genes that are differentially expressed in YFP+ cells compared to YFP-cells. **F-G**) NK cells from c-Myc EGFP reporter mice were either primed prior to stimulation or directly stimulated for 3 hours. EGFP was measured by flow cytometry. **F**) Histogram from a representative experiment. **G**) EGFP quantification shown as mean fluorescence intensity of the stimulated condition relative to their respective unstimulated. n=15 mice, 5 independent experiments.

While *Ifng* transcription is induced in primed NK cells, only ∼50% of cells produce protein (**Fig. 1E**). To more specifically evaluate changes associated with *Ifng* transcription, RNA-seq was performed using primed NK cells from an *Ifng* reporter mouse (GREAT mice, [27]) (**Fig. 5B-C**). These mice serve as a faithful reporter for transcription of *Ifng*, and consistent with our prior data, naïve NK cells from GREAT mice increase YFP expression only after IL-12/18 stimulation, with minimal change with anti-NK1.1 stimulation (**Fig. 5B**). Primed NK cells, however, express YFP after anti-NK1.1 stimulation (**Fig. 5B**). For RNA-seq, primed cells were stimulated with anti-NK1.1 and sorted based on YFP expression. *Ifng* transcript was the most significant gene upregulated in the YFP+ population, demonstrating the utility of this reporter in murine NK cells (**Fig. 5C**). Several pathways were upregulated in the YFP+ (IFN-γ-producing) population compared to the YFP-population, including the unfolded protein response (*Pdia5*, *Stc2*, *Atf3*, *Atf4*, *Atf6*), Myc targets (*Srm*, *Odc1*, *Tfrc*, *Dusp2*), and mTORC1 signaling (*Cth*, *Slc7a11*, *Slc7a5*, *Polr3g*) (**Fig. 5D-E**). Among Myc targets, RNA polymerase III genes *Polr3h* and *Polr3g* were upregulated, while negative regulator of RNA polymerase III *Maf1* was downregulated only in IFN-γ+ cells, supporting that downregulation of this gene is specific to those cells actively transcribing *Ifng* (**Fig. 5E**). Additionally, *Tnf*, *Il23a*, *Spp1*, *Ccl3* and *Xcl1* were upregulated (**Fig. 5E**), indicating that the expression of these genes identified in the previous dataset correlates with *Ifng* transcript expression.

### c-Myc marks NK cells capable of *Ifng* transcription

Myc (c-Myc) is a basic helix-loop-helix transcription factor that forms a heterodimer with Myc associated factor X (Max) to bind E-box sequences and modulate expression of target genes. In NK cells, cytokine (IL-2+IL-12, IL-12+IL-18, or IL-15) stimulation upregulates c-Myc expression [61–63], which, in turn, has been linked to metabolic and effector function changes [61, 64]. Given the convergence of transcriptional evidence here and existing literature suggesting a link between Myc and *Ifng* transcription, we further characterized the potential relationship between Myc and *Ifng* in our system. To assess the expression of c-Myc in NK cells after priming, mice expressing a N-terminal fusion EGFP/MYC protein [28] (c-Myc^EGFP^) were evaluated. IL-12+IL-18 stimulation (3 hours) upregulated Myc protein (marked by GFP expression) in naïve, LD IL-15 cultured, and HD IL-15 primed NK cells. In contrast, ITAM-based anti-NK1.1 stimulation resulted in upregulation of c-Myc only in HD IL-15 primed NK cells (**Fig. 5F, 5G**). Thus, c-Myc expression is upregulated in stimulated NK cells capable of *Ifng* transcription.

One hypothesis is that c-Myc may directly regulate *Ifng* transcription in primed NK cells. The *Ifng* locus features several E-box sequences and degenerate sequences, supporting the possibility of Myc binding and regulating *Ifng* transcription. To determine if Myc transcription factor activity regulates *Ifng* production, a pharmacological approach to inhibit Myc-Max dimerization, and therefore abrogate Myc transcription factor function, was utilized. Primed NK cells were treated with KJ-Pyr-9 [65] during the 6 hour anti-NK1.1 (**Fig. 6A**) or anti-Ly49D stimulation (**Fig. S4A**). KJ-Pyr-9 interferes with Myc-Max complex formation in cells and reduces MYC-driven transcriptional signature [65, 66]. Treatment with KJ-Pyr-9 led to reduced *Ifng* transcript upregulation upon activating receptor stimulation in primed NK cells, and a reversion to metabolic dependence for IFN-γ production (**Fig. 6A-B, S4A-B**). Myc inhibition did not affect cytokine-stimulated *Ifng* transcription, but did inhibit transcription when cytokine-stimulated cells were cultured with the OXPHOS inhibitor oligomycin (**Fig. S4C-D**). This suggests that Myc-Max dimerization may be essential for activating receptor-stimulated *Ifng* transcription, but only becomes important for cytokine-stimulated *Ifng* transcription with metabolic inhibition.

**Fig 6.**
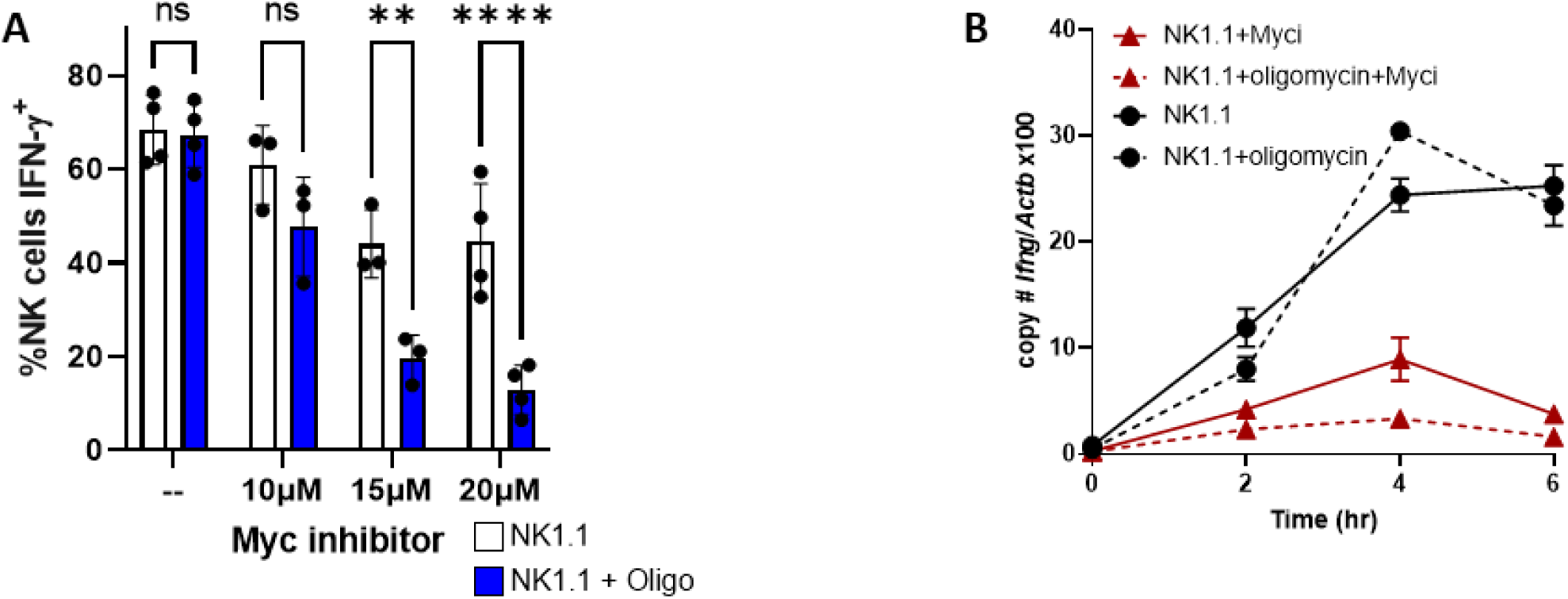
Myc inhibition alters IFN-γ regulation. Purified mouse NK cells were cultured for 72 hours in 100 ng/ml IL-15, washed to remove the cytokine, then stimulated with anti-NK1.1 (**A,B**) +/- Myc inhibitor KJ-Pyr-9 (Myci) +/- OXPHOS inhibitor oligomycin. **A**) % IFN-γ+ NK cells were measured by flow cytometry, n=8 mice, 4 independent experiments; 2way ANOVA, Sidak’s correction for multiple comparison test. **B**) *Ifng* transcript was measured by quantitative RT-PCR normalized to beta-actin (*Actb*). Representative experiment with 1 technical and 1 biological replicate (2 mice), out of 3 independent experiments. Statistical analysis: 2-way ANOVA; * denotes significance from NK1.1 vs. NK1.1+Myci comparison.

Next, a genetic approach in which *Myc* was conditionally deleted in NK cells (*Ncr1*-*Myc*Δ/Δ) *in vitro* with hydroxytamoxifen (4-OHT) was used to determine if Myc is required for *Ifng* upregulation (**Fig. 7A**-**B**). In this model, iCre recombinase nuclear activity was induced by tamoxifen and marked by YFP expression, allowing for detection and temporal control of Myc deletion, without affecting NK development and homeostasis. Enriched NK cells were primed (72 hours, 100 ng/ml IL-15) in the presence of 4-hydroxytamoxifen (48-72 hours, 0.8 uM 4-OHT). Myc deletion was confirmed by real-time PCR comparison of cells marked by Cre activity (YFP+) or not (YFP-) (**Fig. 7B**). In contrast to Myc inhibitor data, there was no significant change in the *Ifng* transcript or the percentage of IFN-γ+ NK cells between YFP+ and YFP-cells after 6 hours of anti-NK1.1 stimulation, regardless of the presence or absence of oligomycin (**Fig. 7C-D**). Furthermore, c-Myc was not required during the IL-15 priming phase for metabolic flexibility for activating receptor stimulated IFN-γ production (**Fig. 7E**). Therefore, c-Myc marks NK cells capable of *Ifng* transcription, but it is not required itself for *Ifng* transcription in IL-15 primed NK cells. There are several possibilities that could explain the contradictory results observed with chemical Myc inhibition, including lack of inhibitor specificity, affecting other leucine zipper-based dimerization of transcription factors, and compensatory activity of another Myc family member in the genetic model.

**Figure 7.**
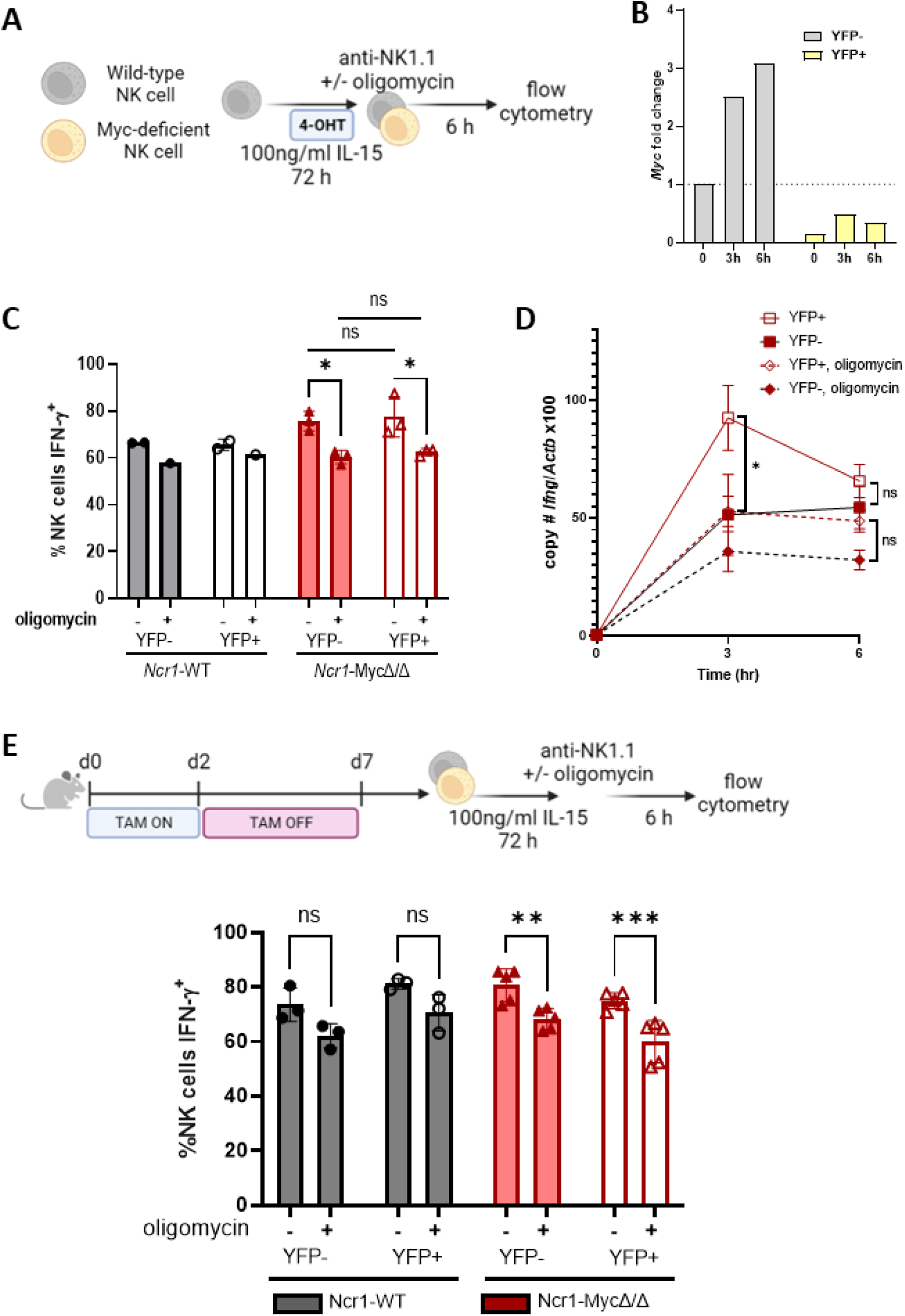
Myc deletion does not affect *Ifng* transcription. **A**) Experimental schema. Enriched splenic NK cells were cultured in 100 ng/ml IL-15. 4-OHT was added after 6-24 hours at a final concentration of 0.8 uM for the remainder of the 72 hours. Cells were washed and then stimulated via plate-bound anti-NK1.1, +/- 100 nM oligomycin. **B**) *Myc* transcript (day 3) normalized to unstimulated YFP- cells from *Ncr1*-MycΔ/Δ; representative experiment. **C**) % IFN-γ+ NK cells; N*cr1*-WT were used as controls. 2 independent experiments. **D**) NK cells from *Ncr1*-MycΔ/Δ mice were primed and stimulated with anti-NK1.1 for up to 6 hours. Absolute copy number of *Ifng* and *Actb* were quantified by real-time PCR for sorted YFP+ and YFP-populations. Results represent the ratio of *Ifng* to *Actb* x 100 and the mean +/- SEM from 3 independent experiments. **E**) Experimental schema and % IFN-γ+ NK cells measured by flow cytometry. 2 independent experiments with replicate wells, 2-6 mice per group. Statistical analysis: 2 way ANOVA.

## DISCUSSION

In this study, we demonstrate that IL-15 priming leads to changes in murine NK regulatory networks, altering IFN-γ regulation and providing metabolic flexibility upon activating receptor stimulation. Such priming has the potential to impart NK cells with enhanced function in metabolically challenging environments such as the tumor microenvironment. Furthermore, IL-15 based therapies have been developed, with improved antitumor activity in preclinical models, especially when combined with immune checkpoint inhibition strategies [22].

Naïve splenic NK cells constitutively express *Ifng* transcript, but not protein, at levels greater than naïve CD4 T cells sorted from lymph nodes in C57BL/6 mice [67]. This suggests a translational repression mechanism occurs in resting murine NK cells. In contrast to IFN-γ regulation in CD4+ T cells, where GAPDH acts as a translation repressor by binding to the *Ifng* 3’UTR [49], we demonstrate here that metabolic regulation of IFN-γ in naïve murine NK cells is 3’UTR-independent. Furthermore, our results suggest the main mode of IFN-γ production in cytokine stimulated naïve NK cells is by production of new *Ifng* transcript, rather than production of the protein from pre-formed *Ifng* transcript, as is the case for activating receptor stimulation.

IL-15 primed NK cells exhibited metabolic flexibility and a switch to transcriptional regulation for activating receptor stimulated IFN-γ production. Glycolysis, and, to a lesser extent, mTORC1, are required during the priming phase for this effect. To investigate mechanisms responsible for the switch to IFN-γ transcriptional regulation in primed NK cells, we tested several hypotheses. ATAC-seq analysis demonstrated that while IL-15 priming leads to broad changes in chromatin accessibility, it does not alter accessibility at the *Ifng* itself. In NK cells, the IFN-γ locus is already accessible and primed for transcription when compared to T cells [67–71]. However, it is possible that methylation and acetylation changes modulate transcription factor occupancy, which is something not explored here. DNA hypomethylation in the *IFNG* locus has been reported during terminal differentiation of human NK cells and in an HCMV adaptive subset with a greater capacity to produce IFN-γ [72, 73].

Canonical ITAM signaling strength was not different between primed and LD IL-15 cultured NK cells for several of the signaling molecules assessed, including MAPK p38, which has been linked to increased *IFNG* transcript stability in human NK cells [74, 75]. However, IL-15 priming led to a new transcriptional profile upon activating receptor stimulation. We hypothesize that changes in chromatin accessibility induced by IL-15 priming fundamentally alter how NK cells receive and respond to signaling downstream from ITAM receptor activation, either directly by chromatin accessibility or potentially through upregulation of pathways that allow for non-canonical signaling. For example, IL-15 priming led to a strong enrichment for AP-1 family motifs in the peaks with increased chromatin accessibility, and these motifs are also enriched in memory (Ly49H+) NK cells in mice [76] and human HCMV adaptive NK cells [77]. This suggests IL-15 induces an epigenetic signature associated with enhanced function.

A significant number of published NK cell studies rely on IL-15 or IL-2, a cytokine with similar effects and shared receptor subunits, to maintain or expand NK cells in culture. High dose IL-15 in vitro significantly alters NK cell metabolism and functional capacity, and, as we show here, fundamentally alters regulation of IFN-γ production [16, 19, 78]. Therefore, it is important to consider how cells are treated ex vivo when trying to extrapolate in vitro findings to what might be occurring in vivo. We demonstrate that NK1.1 activation of IL-15 primed NK cells generated a distinct transcriptional profile compared to LD IL-15 cultured or naïve NK cells and marked a switch to *Ifng* transcriptional regulation. RNA-seq studies identified c-Myc and its targets to be positively correlated with *Ifng* transcription. In T cells, *Myc* is upregulated upon TCR stimulation, and protein levels are then sustained with the help of cytokines (such as IL-2 or IL-15) [79] and amino acid transporter Slc7a5 [80]. Similarly, another study demonstrated that in NK cells, IL-2+IL-12 stimulation leads to c-Myc upregulation, as well as Slc7a5 expression and activity, which in turn is required for c-Myc expression at later time points [61]. Additionally, loss of c-Myc protein *in vitro* reduced the percentage of IFN-γ+ NK cells and the amount of IFN-γ produced per cell after IL-2+IL-12 [61]. While activating receptor stimulation leads to c-Myc upregulation in primed NK cells, loss of c-Myc in these cells did not alter the percentage of IFN-γ+ NK cells with ITAM stimulation. We did observe, however, striking results with Myc:Max heterodimer formation inhibitor KJ-Pyr-9. *Ifng* transcript and the percentage of IFN-γ+ NK cells were reduced, with further reduction in the latter upon OXPHOS inhibition. The differences between the results obtained with the inhibitor and those obtained with the genetic model may simply indicate off-target effects of the inhibitor or a lack of specificity for disruption of Myc:Max dimer formation. Inhibition of oncogenic activity of N-MYC, v-jun and PIK3CA-H1047R (gain of function) has been reported with the inhibitor used in the studies here [65], supporting the possibility of off-target effects. In addition, a separate but not mutually exclusive mechanism may involve other Myc family members compensating for c-Myc’s absence. *Mycn*, encoding for n-Myc, is not normally expressed in NK cells but *Mycn* transcript was upregulated in primed NK cells (**Fig. 4H**). This, together with KJ-Pyr-9 inhibition of N-MYC oncogenic activity [65], offers a potential explanation for the discrepancies between the inhibitor and genetic model results that will require further investigation.

Another pathway observed to be specifically upregulated in IFN-γ+ cells here was the unfolded protein response. This pathway has been linked to production of an active transcriptional activator, spliced XBP1 (XBP1s) [81, 82], which acts downstream of IL-15 to support human NK cell survival [83] and can interact with T-bet to support *GZMB* [83] and *Ifng/IFNG* transcription [83, 84]. Further studies of XBP1s function in NK cells may determine its role in *Ifng* regulation in IL-15 primed NK cells. The UPR pathway has also been linked to *IL23A* regulation and subsequent IL-23 production in myeloid cells [85–87], and may explain the *Il23a* upregulation observed in IFN-γ+ cells.

Overall, this study highlights that IL-15 dose and culture time can dramatically alter chromatin accessibility and regulatory networks in murine NK cells. Functionally, primed NK cells switch to *Ifng* transcriptional regulation in response to ITAM signaling via NK1.1, and no longer require a secondary, metabolism-derived signal for optimal IFN-γ production. Therefore, primed cells have distinct properties from freshly isolated NK cells or NK cells cultured in low dose IL-15, and interpretation of results from NK cell studies should take into account cell culture conditions such as IL-15 dose and time. While IL-15 priming confers this advantage, supporting IL-15 use for immunotherapeutic purposes, IL-15 use must be carefully considered and calibrated in light of reported detrimental effects [22, 88, 89].

## Acknowledgements

The authors would like to thank Kelsey Trammel and Katherine Owens for mouse colony management and technical assistance, and Prabhakar Sairam Andhey for assistance with data processing. We thank Eric Tycksen and the Genome Technology Access Center in the Department of Genetics at Washington University School of Medicine for help with genomic analysis.

**Figures 3A, 4A**, **5C**, **7A**, and part of **7E** were created with BioRender.com.

