## Supplemental figures for "IL-15 priming alters IFN-γ regulation in murine NK cells"

**Figure S1**: Activation-specific requirements for IFN-γ production by human NK cells.

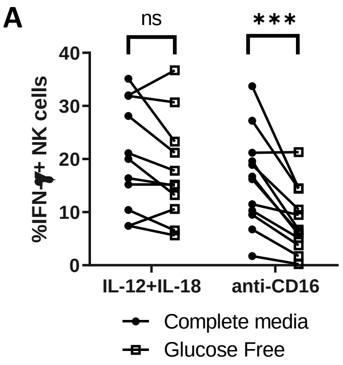


**Figure S1. Activation-specific requirements for IFN-γ production by human NK cells.** Human NK cells stimulated with either IL-12+IL-18 (10 ng/ml and 50 ng/ml, respectively) or with plate-bound anti-CD16 (3G8, 1 mg/ml) in complete or glucose-free media for 6 hours. n=7 donors, 6 independent experiments. Statistical analysis: Paired t-test

**Figure S2**: No IL-15 dose-dependent changes in ITAM signaling with metabolic inhibitors

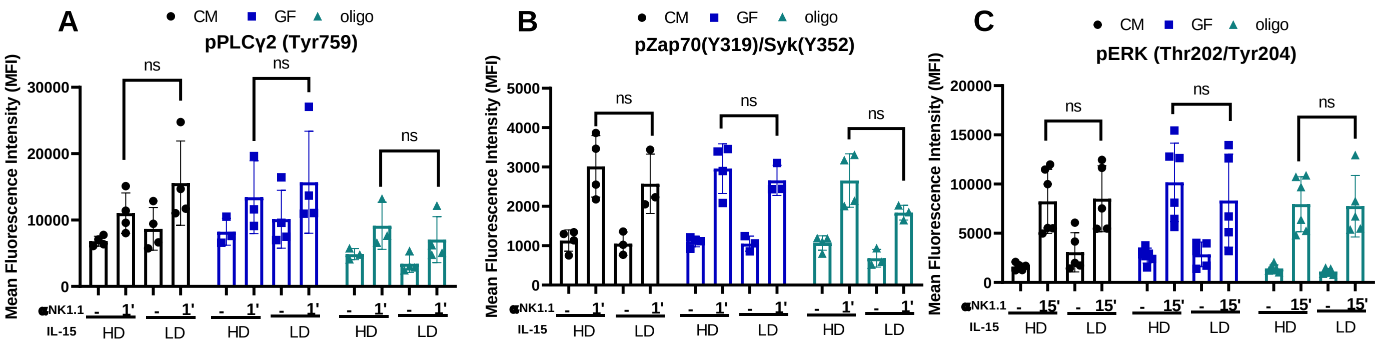


**Figure S2. No IL-15 dose-dependent changes in ITAM signaling with metabolic inhibitors.** NK cells were cultured for 72 hours with 10 ng/ml (LD) or 100 ng/ml (HD) IL-15, followed by anti-NK1.1 stimulation for the indicated time, in complete media (CM), glucose-free media (GF), or complete media + oligomycin (oligo). Phosphorylation of Zap70/SYK (**A**), PLCγ2 (**B**), and ERK (**C**) was quantified by flow cytometry. Results shown are from a minimum of 3 independent experiments, 1-2 mice per experiment. Statistical analysis: Wilcoxon test (paired t-test, non-parametric).

**Figure S3.** RNA-seq supplemental figure

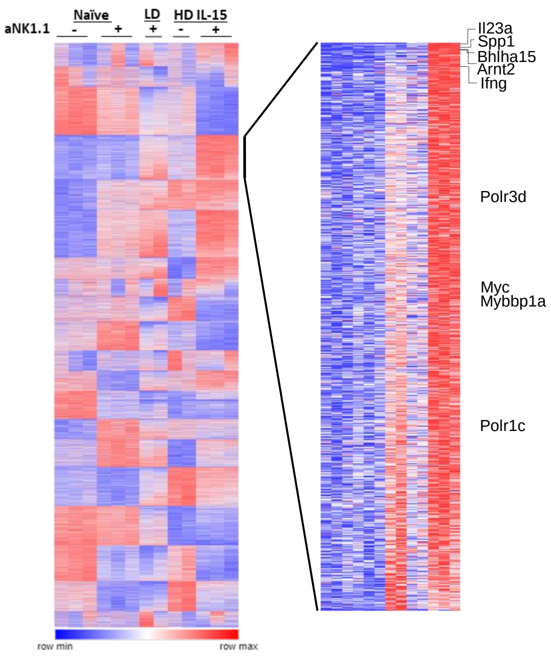


**Figure S3. RNA-seq supplemental figure. A**) k-means clustering strategy highlighting a cluster of genes uniquely upregulated in primed NK1.1-stimulated NK cells. Select genes are annotated.

**Figure S4.** Myc inhibition in murine NK cells.

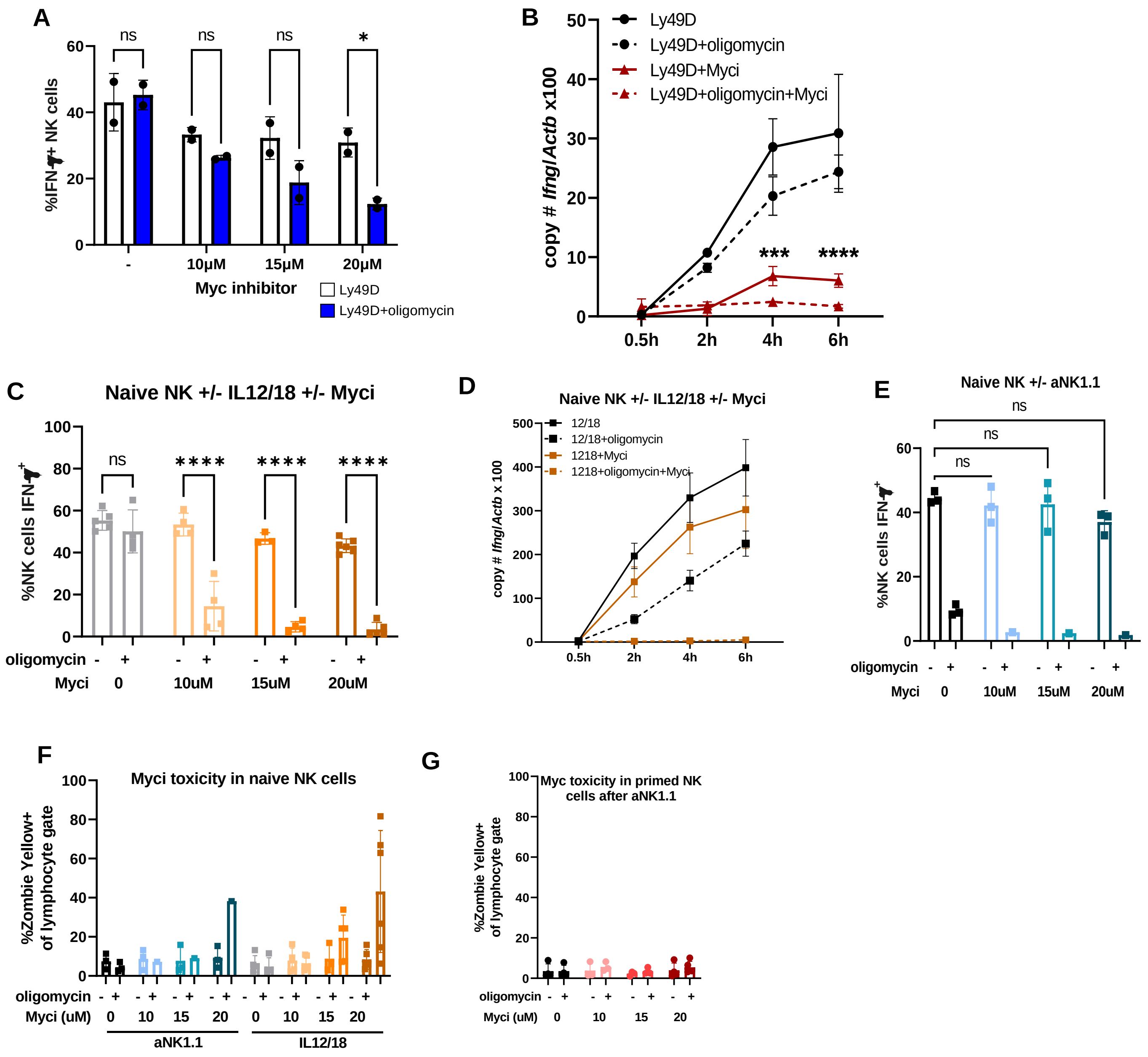


**Fig S4. Myc inhibition in murine NK cells.** Purified mouse NK cells were cultured for 72 hours in 100 ng/ml IL-15, "washed" to remove the cytokine, then stimulated with anti-Ly49D (**A-B**) in the absence or presence of Myc inhibitor KJ-Pyr-9 (Myci) +/- OXPHOS inhibitor oligomycin. n=10 mice, 2 independent experiments. **A**) % IFN-γ+ NK cells were measured by flow cytometry. **B**) *Ifng* transcript was measured by quantitative RT-PCR normalized to beta-actin (*Actb*). * denote significance from NK1.1 vs. NK1.1+Myci comparison. **C-D**) Purified mouse NK cells were stimulated with IL-12 (1ng/ml) and IL-18 (1ng/ml) or anti-NK1.1 for 6 hours +/- of Myc inhibitor KJPyr-9 (Myci) +/- OXPHOS inhibitor oligomycin. **C**) % IFN-γ+ NK cells were measured by flow cytometry after IL12/18 stimulation. 2-4 independent experiments **D**) *Ifng* transcript was measured by quantitative RT-PCR normalized to beta-actin (*Actb*). n=4 independent experiments. * denotes significance from NK1.1 vs. NK1.1+Myci comparison. **E**) % IFN-γ+ NK cells were measured by flow cytometry after anti-NK1.1 stimulation; n=3 independent experiments. Myci toxicity in naïve (**F**) or primed (**G**) NK cells after 6 h stimulation, as measured by Zombie Yellow dye. 3-4 independent experiments. Statistical analysis: 2 way ANOVA.
